# Perturbed fatty-acid metabolism is linked to localized chromatin hyperacetylation, increased stress-response gene expression and resistance to oxidative stress

**DOI:** 10.1101/2022.06.12.495787

**Authors:** Jarmila Princová, Clàudia Salat-Canela, Petr Daněk, Anna Marešová, Laura de Cubas, Jürg Bähler, José Ayté, Elena Hidalgo, Martin Převorovský

**Affiliations:** Laboratory of Microbial Genomics, Department of Cell Biology, Faculty of Science, Charles University, Viničná 7, Prague 2, 128 00, zechia; Oxidative Stress and Cell Cycle Group, Universitat Pompeu Fabra, C/ Dr. Aiguader 88, 08003 Barcelona, Spain; Institute of Healthy Ageing and Department of Genetics, Evolution & Environment, University College London, London WC1E 6BT, UK

## Abstract

Oxidative stress is associated with cardiovascular and neurodegenerative diseases, diabetes, cancer, psychiatric disorders and aging. In order to counteract, eliminate and/or adapt to the sources of stress, cells possess elaborate stress-response mechanisms, which also operate at the level of regulating transcription. Interestingly, it is becoming apparent that the metabolic state of the cell and certain metabolites can directly control the epigenetic information and gene expression. In the fission yeast *Schizosaccharomyces pombe*, the conserved Sty1 stress-activated protein kinase cascade is the main pathway responding to most types of stresses, and regulates the transcription of hundreds of genes via the Atf1 transcription factor. Here we report that fission yeast cells defective in fatty acid synthesis (*cbf11, mga2* and ACC/*cut6* mutants) show increased expression of a subset of stress-response genes. This altered gene expression depends on Sty1, and the Gcn5 and Mst1 histone acetyltransferases, is associated with increased acetylation of histone H3 at lysine 9 in the corresponding gene promoters, and results in increased cellular resistance to oxidative stress. Since both fatty-acid synthesis and histone acetylation compete for the same substrate, acetyl-CoA, we propose that changes in lipid metabolism can regulate the chromatin and transcription of specific stress-response genes, which in turn might help cells to maintain redox homeostasis.

## INTRODUCTION

Oxidative stress occurs when the equilibrium between the production and the detoxification of oxidants is disturbed, leading to damage of cellular molecules (1). Importantly, oxidative stress is associated with multiple cardiovascular and neurodegenerative diseases, diabetes, cancer, psychiatric disorders and aging (2–5). However, under physiological conditions that create increased oxidant levels, such as mitochondrial respiration or fatty acid (FA) oxidation, the cellular antioxidant mechanisms are typically able to maintain cellular redox homeostasis and prevent oxidative stress (1, 6).

In the fission yeast *Schizosaccharomyces pombe*, the Sty1 stress-activated protein kinase (SAPK) cascade, homologous to the mammalian p38 mitogen-activated protein kinase, is the main pathway responding to most types of stress conditions, including oxidative stress. Once activated, Sty1 translocates to the nucleus, where it phosphorylates and activates its main target, the basic zipper-containing transcription factor Atf1, leading to an extensive transcriptional response (7, 8). Analyses of genes showing differential expression in response to various stresses identified the core environmental stress response (CESR) as a group of genes that are jointly regulated under all or most environmental stresses. Remarkably, the regulation of most CESR genes depends on Sty1 and, to a lesser extent, on Atf1 (9, 10).

The CSL (<u>C</u>BF1, <u>S</u>u(H), <u>L</u>ag-1) family protein Cbf11 and the IPT/TIG ankyrin repeat-containing protein Mga2 are transcription factors regulating lipid-metabolism genes. Their target genes include the acetyl-CoA carboxylase *cut6*, the acyl-CoA desaturase *ole1*, the long chain fatty acid-CoA ligases *lcf1* and *lcf2*, or the triacylglycerol lipases *ptl1* and *ptl2* (11, 12). The loss of Mga2 causes a general disruption of the lipidome (12), while cells lacking Cbf11 have a decreased amount of lipid droplets and show mitotic defects (13). Curiously, we have previously shown that many CESR genes are upregulated in the *cbf11Δ* deletion mutant cells, but the reason for these changes is not clear (11).

It has become apparent in recent years that the cellular metabolic state can directly affect the regulation of gene expression through the availability of selected metabolites that serve as substrates for various chromatin modifying enzymes. For example, the metabolite acetyl-CoA is central to multiple biosynthetic pathways as well as to histone acetylation by histone acetyltransferases (HAT_s_). Perturbations in acetyl-CoA levels lead to altered histone acetylation and gene transcription (14–16). Acetyl-CoA is also utilized during FA synthesis, including its first and rate-limiting step catalyzed by the acetyl-CoA carboxylase (ACC). Intriguingly, ACC inhibition increases the acetylation of bulk histones and affects gene expression in yeast (17). However, the significance and the physiological consequences of such interconnections between the metabolic state and gene expression patterns are only beginning to be understood. In this study, we show that a decrease in FA synthesis leads to increased expression of specific stress-response genes accompanied by promoter histone hyperacetylation, and to increased resistance to hydrogen peroxide (H_2_O_2_)-induced oxidative stress in fission yeast.

## MATERIALS AND METHODS

### Plasmid construction

The Cas9/sgRNA_TEFp (pMP134) and Cas9/sgRNA_*cbf11* (pMP153) plasmids were constructed as previously described (18). sgRNAs targeted to the TEF promoter region of the *natMX6* cassette and to *cbf11* ORF, respectively, were inserted into the pMZ374 plasmid carrying an empty sgRNA site and a sequence encoding the Cas9 endonuclease (19). Briefly, the whole pMZ374 was amplified by NEB Q5 polymerase (using AJ11 and AJ12, and AJ29 and AJ30 oligonucleotides, respectively), 5’ ends of purified PCR products were phosphorylated, and plasmid ends were ligated together. The final plasmids were verified by restriction cleavage and sequencing. pMZ374 was a gift from Mikel Zaratiegui (Addgene plasmid # 59896; http://n2t.net/addgene:59896; RRID:Addgene_59896). Lists of oligonucleotides and plasmids used in this study are provided in Supplementary Tables 1 and 2, respectively.

### Strains, media and cultivations

Fission yeast cells were grown according to standard procedures (20) in either complex yeast extract medium with supplements (YES) or Edinburgh minimal medium (EMM). A list of strains used in this study is provided in Supplementary Table 3.

For construction of Cbf11-TAP scarless knock-in strain, the CRISPR/Cas9-based strategy was adapted from (18). MP15 cells (*h-cbf11-ctap4::natR ura4-D18 leu1-32 ade6-M216*) (11) were synchronized in G1 and transformed with a fragment of the *cbf11-TAP* sequence (plasmid pMaP27 digested by SalI and EcoO109I) as template for homologous recombination together with the Cas9/sgRNA_*TEFp* plasmid (pMP134). After selection on EMM+ade+leu plates, the smallest colonies were re-streaked onto non-selective YES plates to allow for elimination of the deleterious Cas9 plasmid. The integration of *cbf11-TAP* was verified by PCR (primers MaP169 and MP28) and sequencing. Expression of Cbf11-TAP protein was verified by western blot with an anti-TAP antibody (Thermo Scientific, CAB1001). Prototrophic *cbf11-TAP* strain was then prepared by standard crossing and revalidated.

The Cbf11DBM-TAP scarless knock-in strain was constructed and validated analogously in two steps. First, to insert the DNA-binding mutation (R318H; DBM) (21) into the *cbf11* endogenous locus MaP70 cells (*h-cbf11-3HA::natMX6 ura4-D18 leu1-32 ade6-M216*) were synchronized in G1 phase and transformed with a Cas9/sgRNA plasmid targeting Cas9 next to the desired DBM mutation site in *cbf11* ORF (Cas9/sgRNA_*cbf11;* pMP153), and a *cbf11DBM-TAP* DNA fragment as template for homologous recombination (plasmid pMaP11 digested by SalI and EcoO109I). Introduction of the DBM mutation was verified by PCR (primers MP53 and MP54) coupled with restriction digestion with HpaII, and by sequencing. The resulting strain (MP670) contained the DBM mutation, but retained the HA tag and *natMX6* cassette at the *cbf11* locus. In the second step the MP670 cells were transformed with the pMP134 plasmid targeting the *natMX6* cassette, and the pMaP11 fragment described above as template for homologous recombination. The final prototrophic strain *cbf11DBM-TAP* (MP712) was then prepared by standard crossing and validated by PCR, restriction cleavage, sequencing and western blot as described above.

All other strains constructed in this study were created using standard genetic methods (22), or the pClone system (23) for deletion of *cbf11, ssp2* or *mga2* genes.

### Spot tests

Exponentially growing cells were 10-fold serially diluted and spotted onto YES plates containing various concentrations of H_2_O_2_ (Sigma-Aldrich, H1009) or 125 μM menadione sodium bisulfite (Sigma-Aldrich, M5750) for oxidative stress-resistance assays, or 3-MB-PP1 (Sigma-Aldrich, 529582) for Sty1 inhibition. The spots were allowed to dry and plates were incubated at 30°C or 32°C until cell growth was evident. Due to the unstable nature of H_2_O_2_, the plates were poured on the day of spotting and YES agar was cooled to 45°C prior to adding the stressor.

### RT-qPCR

The RT-qPCR results shown in Fig 1B (bottom panel), Fig 2A, B, Fig 4D, Fig 5B-D and Fig EV5 were obtained as follows: Total RNA was extracted from cells using the MasterPure Yeast RNA Purification Kit including a DNase treatment step (Epicentre), and converted to cDNA using random primers and the RevertAid Reverse Transcriptase kit (ThermoFisher Scientific). Quantitative PCR was performed using the 5x HOT FIREPol® EvaGreen® qPCR Supermix (Solis Biodyne) and the LightCycler 480 II instrument (Roche). For RT-qPCR, *act1* (actin) and *rho1* (Rho1 GTPase) were used as reference genes. The primers used are listed in Supplementary Table 1.

**Figure 1.**
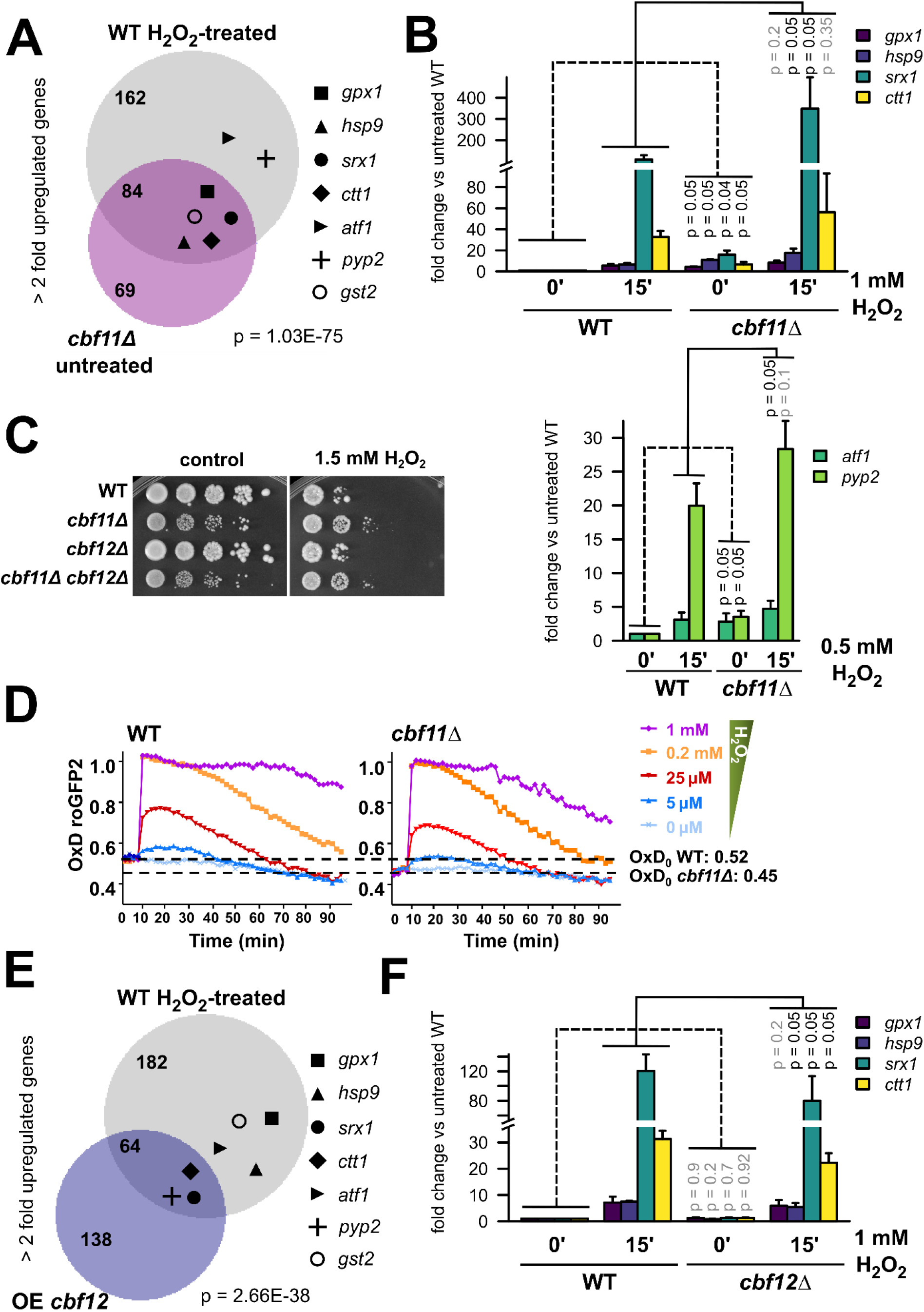
Absence of Cbf11 leads to activation of stress-response genes. **(A)** Venn diagram of genes upregulated more than 2-fold in WT cells after 15 min treatment with 1 mM H_2_O_2_ in EMM medium (data from (29)), and genes upregulated more than 2-fold in *cbf11Δ* cells growing exponentially in YES medium (data from (11)). The group membership of stress genes from panel B is indicated with symbols. Overlap significance was determined by two-sided Fisher’s exact test. **(B)** Expression of the indicated stress genes in WT and *cbf11Δ* cells treated or not with the indicated concentrations of H_2_O_2_ for 15 min in YES was analyzed by RT-qPCR. Mean and SD values of three independent replicates are shown. One-sided Mann-Whitney U test was used to determine statistical significance. **(C)** Survival and growth under oxidative stress of WT, *cbf11Δ, cbf12Δ* and *cbf11Δ cbf12Δ* cultures spotted on YES plates containing 1.5 mM H_2_O_2_. **(D)** *cbf11Δ* displays lower steady-state intracellular H_2_O_2_ levels, and detoxifies extracellular peroxides faster than WT cells. The indicated concentrations of H_2_O_2_ were directly added to EMM cultures of WT and *cbf11Δ* cells transformed with plasmid p407.C169S. The degree of probe oxidation on scale from 0 (fully reduced) to 1 (fully oxidized) is indicated on the Y axis (OxD roGFP2). The starting levels of probe oxidation in each strain background (OxD_0_) are indicated by dashed horizontal lines. Mean values from three biological replicates are shown. **(E)** Venn diagram of genes upregulated more than 2-fold in WT cells after 15 min treatment with 1 mM H_2_O_2_ in EMM medium (data from (29)), and genes upregulated more than 2-fold in cells overexpressing Cbf12 grown in EMM (data from (11)). The group membership of stress genes from panel B is indicated with symbols. Overlap significance was determined by two-sided Fisher’s exact test. **(F)** Expression of the indicated stress genes in WT and *cbf12Δ* cells treated or not with 1 mM H_2_O_2_ for 15 min in YES was analyzed by RT-qPCR. Mean and SD values of three independent replicates are shown. One-sided Mann-Whitney U test was used to determine statistical significance.

**Figure 2.**
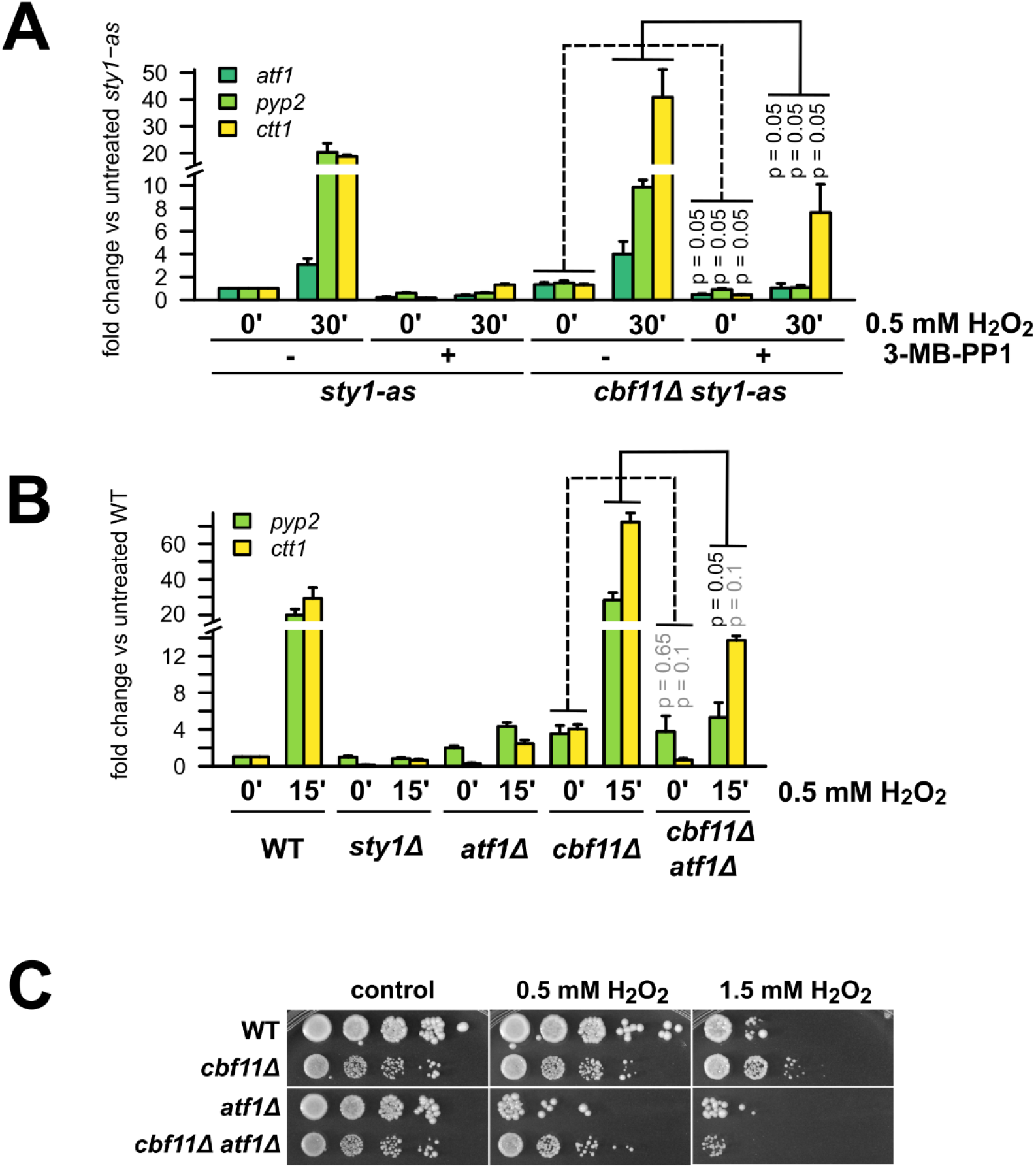
Stress-gene activation in *cbf11Δ* cells is dependent on Sty1 and Atf1. **(A)** Expression of the indicated stress genes in *sty1-as* and *cbf11Δ sty1-as* cells treated or not with the Sty1-as inhibitor 3-MB-PP1 and 0.5 mM H_2_O_2_ for 30 min in YES medium was analyzed by RT-qPCR. Mean and SD values of three independent replicates are shown. One-sided Mann-Whitney U test was used to determine statistical significance. **(B)** Expression of the indicated stress genes in WT, *sty1Δ, atf1Δ, cbf11Δ*, and *cbf11Δ atf1Δ* cells treated or not with 0.5 mM H_2_O_2_ for 15 min in YES was analyzed by RT-qPCR. Mean and SD values of three independent replicates for *pyp2* and two independent replicates for *ctt1* transcript are shown. One-sided Mann-Whitney U test was used to determine statistical significance. **(C)** Survival and growth under oxidative stress of WT, *cbf11Δ, atf1Δ* and *cbf11Δ atf1Δ* cultures spotted on YES plates containing the indicated concentrations of H_2_O_2_.

The RT-qPCR results shown in Fig 1B (top panel), F were obtained as follows: Total RNA was extracted from 40 ml of cells at logarithmic phase by standard phenol/chloroform method, as described earlier (24). 100 μg of total RNA was incubated with recombinant DNase I (Roche, 04716728001) for 30 min at 37°C and the reaction was stopped by incubation at 75°C for 10 min. Reverse transcription was performed on 2 μg of DNase-treated RNA using High-Capacity cDNA Reverse Transcription Kit (Applied Biosystems, 00777852) following manufacturer’s instructions. The cDNA was diluted 1:2 prior to PCR amplification. cDNA was quantified by Real-Time PCR on Light Cycler II using Light Cycler 480 SYBR Green I Master (Roche, 04887352001).

One-sided Mann-Whitney U test was used to determine statistical significance. All statistical tests were performed on data normalized only to reference genes, however, gene expression values normalized to WT or other suitable control sample were used for plotting to allow for more intelligible visualization.

### *In vivo* measurement of roGFP2-Tpx1.C169S oxidation

Wild-type (WT) and *cbf11Δ* cells were transformed with plasmid p407.C169S, constitutively expressing the H_2_O_2_ reporter protein roGFP2-Tpx1.C169S. To measure basal and induced H_2_O_2_ levels, fluorescence of the probe was determined as described before (25). Briefly, roGFP2 exhibits two excitation maxima at 400 nm and 475–490 nm when fluorescence emission is monitored at 510 nm, and the ratio between the two maxima varies upon oxidation by peroxides. Strains were grown in EMM to an OD_600_ of 1, and fluorescence of the cultures was analyzed in 96-well plates before and after the addition of extracellular H_2_O_2_. For calculation of the degree of oxidation of the sensor (OxD), we first subtracted the equivalent fluorescence values of cells lacking the plasmid, and then used the formula displayed in (25).

### TCA extracts and immunoblotting

TCA extracts were prepared as described (26). Atf1 was detected with a polyclonal anti-Atf1 antibody (27), phosphorylated Sty1 was detected with an anti-phospho-p38 antibody (Cell Signaling, 9215), and total Sty1 protein was detected with a polyclonal antibody (28).

### Microarray analysis

Cells were grown to exponential phase (OD_600_ 0.5) in the YES medium at 32°C. At time 0, H_2_O_2_ was added to all cultures to the final concentration of 0.74 mM. Culture aliquots were harvested immediately before, and 15 and 60 minutes after H_2_O_2_ addition by centrifugation for 2 minutes at 1000 g, room temperature, and then snap frozen in liquid nitrogen. RNA extraction and labelling, sample hybridization to custom in-house dual-colour cDNA microarrays, and microarray data processing were performed as described previously (11). Individual Cy3-labelled samples were hybridized together with a Cy5-labelled reference pool (an equimolar pool from all 9 samples in this study). The microarray data are available from the ArrayExpress database (https://www.ebi.ac.uk/arrayexpress/) under accession number E-MTAB-6761.

### ChIP-qPCR

ChIP-qPCR analysis of Cbf11-TAP shown in Fig 3C was performed as described previously (11).

**Figure 3.**
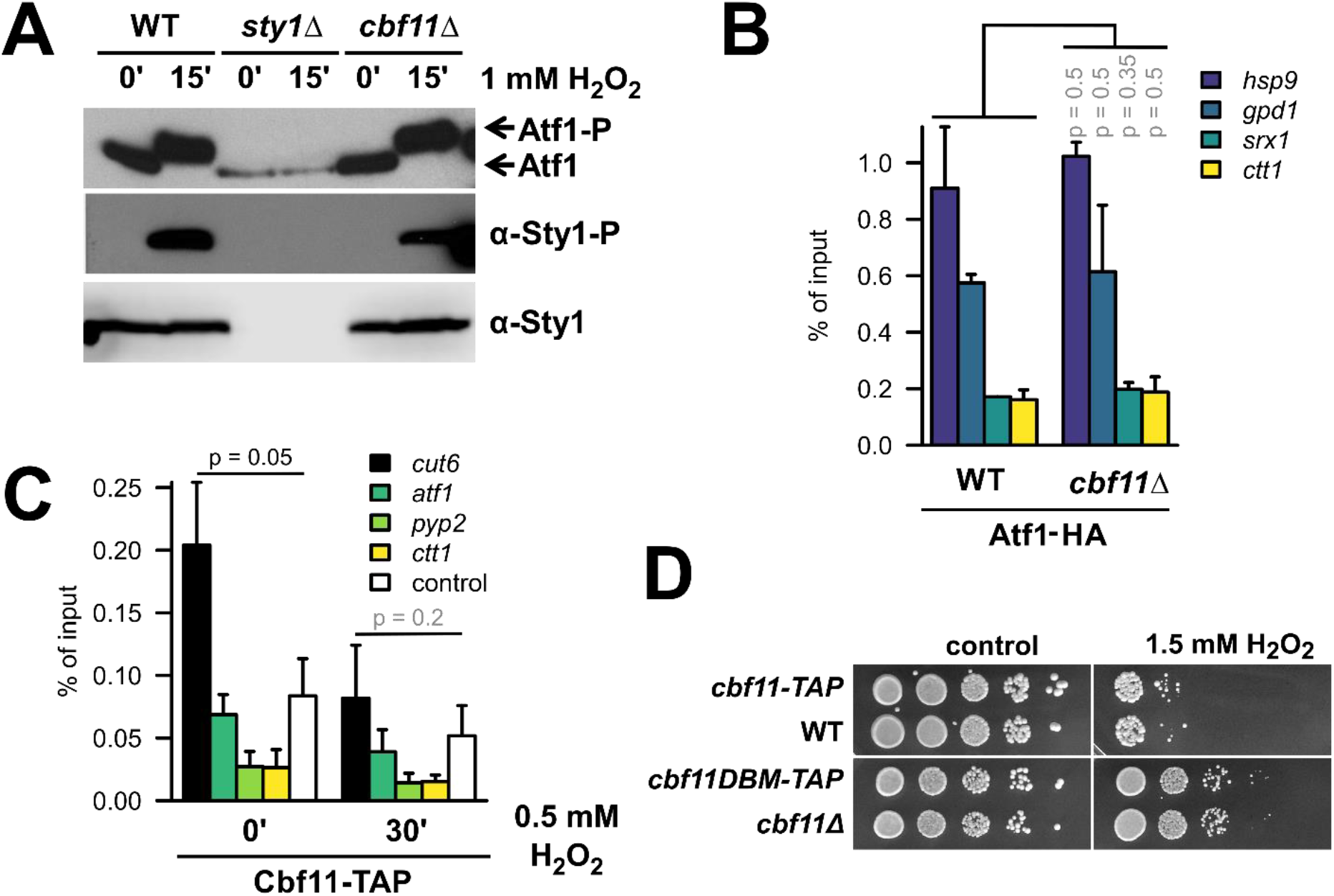
Cbf11 likely affects stress-gene expression indirectly. **(A)** Western blot analysis of Atf1 and Sty1 phosphorylation in WT, *sty1Δ*, and *cbf11Δ* cells treated or not with 1 mM H_2_O_2_ for 15 min in YES medium. Atf1 phosphorylation manifests as retarded migration through the gel. **(B)** Recruitment of Atf1 to the indicated stress-gene promoters was analyzed by ChIP-qPCR in untreated cells grown in EMM medium. Mean and SD values of two independent replicates are shown. Two-sided Mann-Whitney U test was used to determine statistical significance. **(C)** Recruitment of Cbf11 to the indicated stress-gene promoters was analyzed by ChIP-qPCR in cells treated or not with 0.5 mM H_2_O_2_ for 30 min in YES. The *cut6* promoter is a positive control for Cbf11 binding (13); ”control” is an intergenic locus with no Cbf11 binding (Chr I: 1,928,359-1,928,274). Mean and SD values of three independent replicates are shown. One-sided Mann-Whitney U test was used to determine statistical significance; since no stress promoter loci showed signal higher than the negative control locus, these loci were not tested statistically. **(D)** Survival and growth under oxidative stress of WT, *cbf11Δ, cbf11-TAP*, and *cbf11DBM-TAP* cultures spotted on YES plates containing 1.5 mM H_2_O_2_.

For ChIP-qPCR shown in Fig 3B and Fig EV3A, B cells were grown in EMM medium and chromatin isolation was performed as described elsewhere (29). Commercial IgG Sepharose Beads (GE Healthcare, 17-0969-01) were used in order to immunoprecipitate TAP-tagged proteins.

### H3 and H3K9ac ChIP-seq

Two independent replicates were performed. Cells were cultivated to exponential growth phase (OD_600_ 0.5) in the complex YES medium and fixed by adding formaldehyde to the final concentration of 1%. After 30 min incubation, the remaining formaldehyde was quenched by 125 mM glycine. Cells were washed with PBS and broken with glass beads. Extracted chromatin was sheared with the Bioruptor sonicator (Diagenode) using 15 or 30 cycles (for biological replicate 1 and 2, respectively) of 30 s on, 30 s off at high power settings. For all immunoprecipitations (IP) within a biological replicate the same amount of chromatin extract was used (2.5 or 3.7 mg of total protein); 1/10 of the total chromatin extract amount was kept for input DNA control. For each IP 5 μg of antibody (H3: Ab1791, H3K9ac: Ab4441, all Abcam) were incubated with the chromatin extract for 1 hour at 4°C with rotation. Then, 50 μl of BSA-blocked Protein A-coated magnetic beads (ThermoFisherScientific, 10002D) were added to the chromatin extract-antibody suspension and incubated for further 4 hours at 4°C with rotation. The precipitated material and input chromatin extract were decrosslinked, treated with RNase A and proteinase K. DNA was purified using phenol-chloroform extraction and sodium acetate/ethanol precipitation. In biological replicate 2, DNA purification on AMPure XP beads (Beckman Coulter, AC63880) was performed following the phenol-chloroform extraction to remove low-molecular fragments and RNA. Concentration of DNA was measured using the Quantus fluorometer (Promega) and fragment size distribution was checked on Agilent Bioanalyzer using the High Sensitivity DNA Assay. Library construction and sequencing were performed by BGI Tech Solutions (Hong Kong) using the BGISEQ-500 sequencing system.

### ChIP-seq data analysis

The *S. pombe* reference genome sequence and annotation were obtained from PomBase (release date 2018-09-04) (30, 31). Read quality was checked using FastQC version 0.11.8 (https://www.bioinformatics.babraham.ac.uk/projects/fastqc/), and reads were aligned to the *S. pombe* genome using HISAT2 2.1.0 (32) and SAMtools 1.9 (33, 34). Read coverage tracks (i.e., target protein occupancy) were then computed and normalized to the respective mapped library sizes using deepTools 3.3.1 (35). The list of genes upregulated as part of the core environmental stress response (“CESR-UP” genes) was obtained from (9). The CESR-UP genes were further divided into two groups based on whether or not the genes were also upregulated in untreated *cbf11Δ* cells (11). The deepTools 3.5.1 were then used to create average-gene H3 and H3K9ac occupancy profiles for the respective CESR-UP gene subgroups and for all fission yeast genes as a control. The raw ChIP-seq data are available from the ArrayExpress database under the accession number E-MTAB-11081. The scripts used for ChIP-seq data processing and analysis are available from https://github.com/mprevorovsky/ox-stress_histones.

### Acetyl-CoA measurement

Cells grown to exponential phase (OD_600_ 0.5) in the YES medium were collected by centrifugation (1000 g, 5 min) or vacuum filtration (25 ODs). The filter with cell pellet was immediately transferred to 25 ml methanol (−20°C). Samples were centrifuged (3000 g, 5 min, - 4°C, brake 5) and supernatant was decanted. Then 410 μl of 50% freezer-cooled methanol with 10 μM PIPES was added. Cells were broken using glass beads on FastPrep 6.5 m/s, 20 s, 6 cycles. Crude extracts were ultrafiltered using 10kDa filter Amicon Ultra 0.5 ml Ultracel-10K (UFC501096). Samples were evaporated at room temperature on SpeedVac and dissolved in 40 μl 50% acetonitrile. The samples were analyzed on a Dionex Ultimate 3000RS liquid chromatography system coupled to a TSQ Quantiva mass spectrometer (ThermoScientific). A ZIC®-HILIC column (150 mm × 2.1 mm, 5 μm, Merck) was used for separation of analytes. The column was maintained at room temperature and an injection of 1-2 μl of the sample was applied. The gradient elution took 20.5 min and was set from 5% A to 70% A and then 70% A was held for 2 min. A column equilibration step followed and lasted 9 min (A: 10 mM ammonium bicarbonate pH 9.3, B: 97% acetonitrile, flow rate 200 μl/min). Electrospray ionization with switching polarity mode ran under following conditions: ion transfer tube temperature 350 °C, vaporizer temperature 275°C, spray voltage 3500/3000 V (depends on the polarity mode), sheath gas 35 and aux gas 15. For targeted determination of analytes, SRM assay was developed previously by infusing pure compounds.

### Growth curves

OD_600_ was recorded during 24-30 h for cells growing in YE at 30°C from an initial OD_600_ of 0.1 using an automated measurement as previously described (36). When indicated, H_2_O_2_ was added to the cultures.

## RESULTS

### Absence of Cbf11 leads to upregulation of stress-response genes and increased resistance to H_2_O_2_

In our previous studies, we identified fission yeast genes that change their expression upon genetic manipulation of the *cbf11* and/or *cbf12* CSL transcription factor genes (deletion, overexpression). We also noted that these deregulated genes were enriched for stress-response genes (11). More recently, we described the transcriptional signatures of wild-type (WT) cells under oxidative stress (29). Interestingly, when comparing these datasets, we found that 55% of the genes upregulated more than two-fold in cells lacking Cbf11 were also upregulated under oxidative stress in a WT strain (p = 1.03 × 10^−75^; Fig 1A). This finding raises the possibility that a genuine oxidative-stress response is triggered in cells lacking Cbf11, and/or that Cbf11 acts as a direct or indirect repressor of oxidative stress-response genes.

To examine these possibilities further, we first validated our genome-wide data using RT-qPCR. To this end, we selected representative stress-response genes: *srx1, ctt1, gpx1* and *hsp9*, which code for sulfiredoxin, catalase, glutathione peroxidase, and heat shock protein 9, respectively (29). We also included *atf1* and *pyp2*, which encode a transcription factor and a protein phosphatase that regulate the cellular response to oxidative stress (8, 37), and were previously reported to be upregulated ~1.8-fold in *cbf11Δ* (11). We found that all selected genes indeed showed moderately increased basal expression in untreated *cbf11Δ* compared to WT, and became further upregulated upon H_2_O_2_ treatment, reaching even higher transcript levels than in WT (Fig 1B).

Next, we tested whether the upregulation of oxidative stress-response genes in *cbf11Δ* cells has any physiological consequences. We found that compared to WT, *cbf11Δ* cells were more resistant to H_2_O_2_, both when grown on solid media (Fig 1C) and in liquid cultures (Fig EV1A-D). Furthermore, when we introduced the roGFP2-Tpx1.C169S peroxide-sensitive redox probe (25), *cbf11Δ* cells showed lower basal levels of probe oxidation (OxD_0_ of 0.45), which indicates lower steady-state levels of intracellular H_2_O_2_ compared to WT (OxD_0_ of 0.52) (Fig 1D). Of note, a decrease of the OxD_0_ for roGFP2-Tpx1.C169S has previously been reported for cells expressing constitutively active Sty1 (25). Furthermore, cells lacking Cbf11 were also able to detoxify extracellular peroxides faster than WT (compare the reduction slopes of WT and *cbf11Δ* cells in Fig 1D), suggesting a higher peroxide scavenging capacity of these cells. Notably, *cbf11Δ* cells are not resistant to the superoxide generator menadione (Fig EV1E), which triggers a different type of stress response than H_2_O_2_ does (10, 38). Moreover, *cbf11Δ* cells are sensitive to cold stress (39), hypoxia (12) and the microtubule poison thiabendazole (40), suggesting their resistance to H_2_O_2_ is a highly specific phenomenon. Taken together, we have confirmed that oxidative stress-response genes are moderately upregulated in untreated *cbf11Δ* cells, and these cells are resistant to oxidative stress triggered by H_2_O_2_.

We have previously shown that Cbf12, the other fission yeast CSL paralog, acts as a Cbf11 antagonist (11, 39). Therefore, we investigated whether Cbf12 could positively regulate the transcriptional response to oxidative stress. We found that ~32% of the genes upregulated in cells overexpressing Cbf12 were also induced after oxidative stress treatment in WT cells (p = 2.66 × 10^−38^), even though only half of our selected reference genes (*ctt1, srx1* and *pyp2*), were upregulated more than two-fold under both conditions (Fig 1E). However, when we analyzed *cbf12Δ* cells using RT-qPCR, we did not observe any decrease in basal transcript levels of the selected stress-response genes, and we only detected a slightly weaker stress gene induction after stress imposition compared to WT (Fig 1F). Furthermore, we did not detect any notable impact of loss of Cbf12 on the transcriptome-wide response to oxidative stress (see below, Fig 4A). Finally, the *cbf12Δ* strain did not show altered resistance to H_2_O_2_ (Fig 1C) or menadione (Fig EV1E), indicating that Cbf12 only plays a minor role, if any, in the cellular response to oxidative stress. For that reason, we decided to focus on the role of Cbf11 in modulating stress-gene expression.

**Figure 4.**
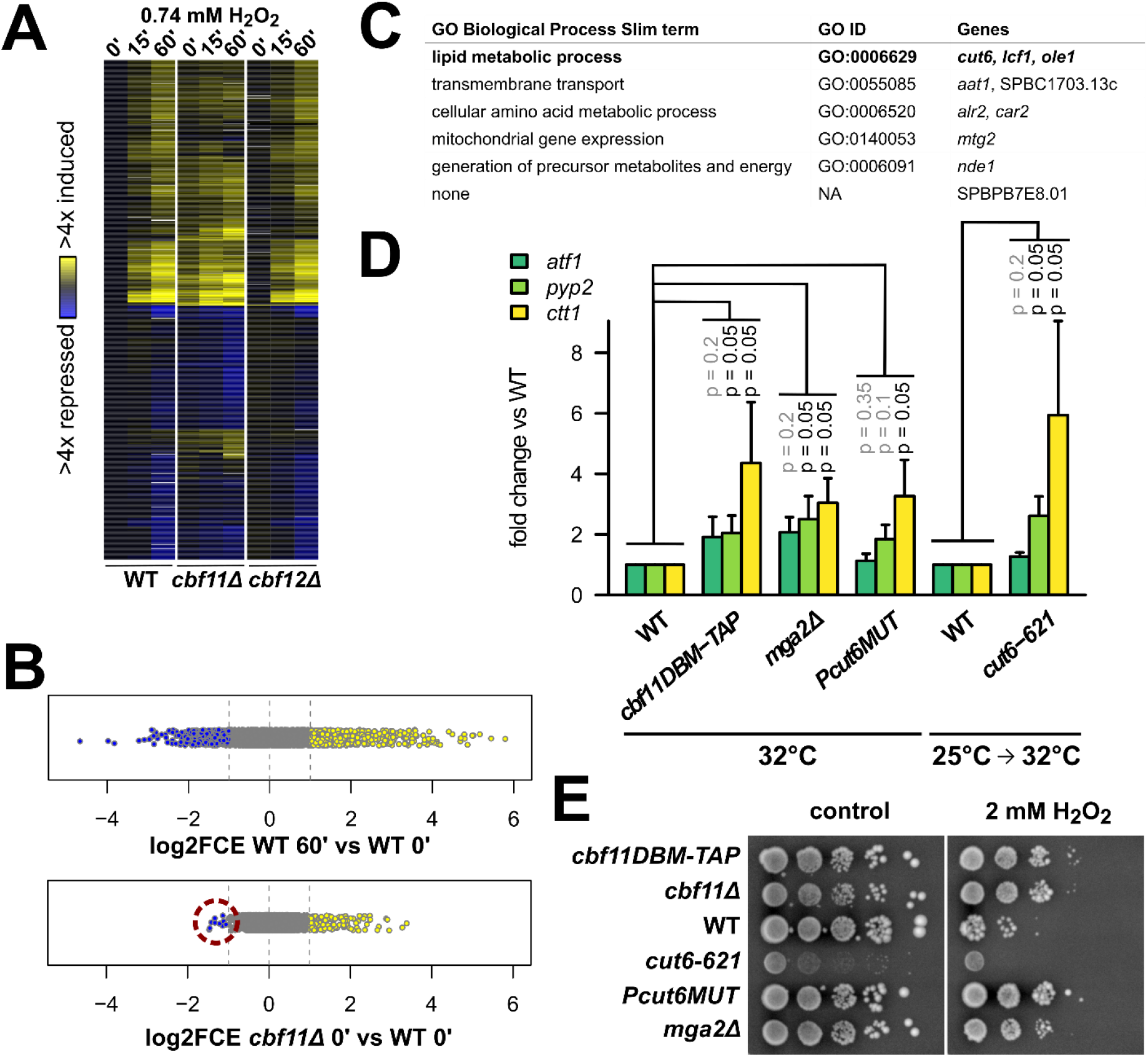
Perturbation of lipid metabolism leads to derepression of stress genes. **(A)** Heatmap of gene expression in WT, *cbf11Δ*, and *cbf12Δ* cells upon treatment with 0.74 mM H_2_O_2_ in YES medium. Transcript levels were determined by a timecourse microarray analysis and normalized to untreated WT values. Only genes showing ≥2-fold change in expression in at least one condition are shown. This experiment was performed once. **(B)** Distribution of all relative transcript levels in WT cells treated for 60 min with H_2_O_2_ and untreated *cbf11Δ* cells from the experiment in panel A. Values represent log2 of fold-changes in expression compared to untreated WT. Individual genes are shown as dots; blue and yellow dots represent ≥2-fold down- and upregulated genes, respectively. The 10 genes downregulated in untreated *cbf11Δ* cells are highlighted with a red circle. **(C)** Biological functions of the 10 genes downregulated in untreated *cbf11Δ* cells from panel B were determined using slimmed Gene Ontology Biological Process annotations (30). Known direct Cbf11 targets are in bold. **(D)** Expression of the indicated stress genes in untreated WT, *cbf11DBM-TAP*, *mga2Δ, Pcut6MUT*, and *cut6-621* cells growing in YES was analyzed by RT-qPCR. The temperature sensitive *cut6-621* mutant was shifted to a semi-restrictive temperature to inhibit Cut6 function prior to analysis. Mean and SD values of three independent replicates are shown. One-sided Mann-Whitney U test was used to determine statistical significance. **(E)** Survival and growth under oxidative stress of WT, *cbf11Δ, cbf11DBM-TAP*, *mga2Δ, Pcut6MUT*, and *cut6-621* cultures spotted on YES plates containing 2 mM H_2_O_2_. The plates were incubated at 32°C, which is a semi-restrictive temperature for the *cut6-621* mutant.

### Increased stress-gene expression in *cbf11Δ* cells depends on the Sty1-Atf1 pathway

Expression of oxidative stress-response genes and resistance to H_2_O_2_ in WT fission yeast cells are critically dependent on the Sty1 kinase and its downstream target, the Atf1 transcription factor (10, 41). Therefore, we tested the requirement for Sty1 and Atf1 in the oxidative stress-related phenotypes of *cbf11Δ* cells.

We previously failed to construct the *cbf11Δ sty1Δ* strain and noted a strong selective pressure for retaining a translocated copy of the *cbf11* gene, suggesting that the double mutant is very sick or lethal (11). Cbf11 and Sty1 exert antagonistic effects on cell-cycle progression, where the G2/M transition is accelerated in *cbf11Δ* cells and delayed in *sty1Δ* cells. Therefore, the poor viability of *cbf11Δ sty1Δ* cells might be due to a conflict in cell-cycle regulation (11, 42, 43). To circumvent these issues, we used an analog-sensitive Sty1 allele in the current study (44). As expected, the *cbf11Δ sty1-as* double mutant showed impaired growth upon Sty1 inhibition (Fig EV2).

Next, we determined whether Sty1 kinase activity is needed for triggering the transcriptional response to H_2_O_2_ in *cbf11Δ* cells. Inhibition of Sty1-as indeed led to severely reduced induction of stress-gene transcripts in H_2_O_2_ in both WT and *cbf11Δ* (Fig 2A, timepoints 30’). Importantly, even the basal transcript levels of oxidative stress-response genes in untreated *cbf11Δ* cells were partially dependent on the Sty1 kinase activity (Fig 2A, timepoints 0’). Similarly, expression of stress-response genes in *cbf11Δ* cells was also partially dependent on Atf1 (Fig 2B), even though the impact of *atf1* deletion was not as severe as Sty1 inhibition. Furthermore, we found that Atf1 was also partially required for the increased resistance of *cbf11Δ* cells to H_2_O_2_ both on solid media and in liquid culture (Fig 2C, Fig EV1B). In summary, these data suggest that the Sty1-Atf1 pathway is required for the moderate upregulation of stress-response genes observed in untreated *cbf11Δ* cells, and for the increased tolerance to H_2_O_2_ of the *cbf11Δ* strain.

### The impact of Cbf11 on stress-gene expression is likely indirect

Since the canonical Sty1-Atf1 stress-response pathway is required for stress-gene upregulation in *cbf11Δ* cells (Fig 2), we explored the following possibilities: 1) *cbf11Δ* cells might experience intrinsic oxidative stress that would result in the activation of the Sty1-Atf1 pathway, or 2) Cbf11 might counteract the activation of the Sty1-Atf1 cascade in WT cells. To test these hypotheses, we first analyzed the phosphorylation (i.e., activation) status of both Sty1 and Atf1 in cells lacking Cbf11. As shown in Fig 3A, no increase in phosphorylation of either Sty1 or Atf1 was observed in *cbf11Δ* cells, suggesting that the canonical stress response is not triggered in unstressed *cbf11Δ* cells, and neither does Cbf11 seem to block Sty1 activation.

Next, we tested whether Cbf11 could block the binding of Atf1 to stress-responsive promoters. We have previously reported two different subsets of stress genes: (i) in unstressed conditions, Atf1 is pre-bound to the first subset of genes, including *gpd1* and *hsp9*, and it is not recruited further after stress imposition; (ii) without stress, Atf1 shows relatively low occupancy at the second subset of genes, which includes *ctt1* and *srx1*, but is further recruited after stress imposition due to the activation of other transcription factors such as Pap1 (29). We found that Atf1 promoter occupancy in unstressed cells was not altered in *cbf11Δ* compared to WT (Fig 3B). Therefore, it is unlikely that Cbf11 could block or compete with Atf1 for binding to stress gene promoters.

Since Cbf11 was previously shown to co-precipitate with Atf1 (45), we also examined the possibility of Cbf11 recruitment to the promoters of stress genes, where it could directly repress their expression under non-stressed conditions, although our previous Cbf11 ChIP-seq analysis did not indicate a clear presence of Cbf11 at stress genes (11). Nevertheless, we tried to detect Cbf11 by ChIP-qPCR at several stress-gene promoters (both in untreated and H_2_O_2_-treated cells), focusing on potential (weak) Cbf11 ChIP-seq peaks (Fig 3C) and known Atf1 binding sites (Fig EV3). The promoter of *cut6*, a well-characterized Cbf11 target gene involved in fatty-acid synthesis, served as a positive control. We assayed Cbf11 binding to DNA in both YES and EMM media, using HA- and TAP-tagged Cbf11, but we did not detect clear Cbf11 presence at the tested stress genes. Collectively, these results strongly argue that the repressive effect of Cbf11 on Sty1/Atf1-dependent stress-gene expression is brought about indirectly.

The R318H substitution in the beta-trefoil domain of Cbf11 (Cbf11DBM) abolishes its binding to the canonical CSL response element (21). To test whether its DNA-binding activity is at all required for Cbf11 to repress Sty1-Atf1 target genes, we introduced the DBM mutation into the endogenous *cbf11* locus. Notably, the *cbf11DBM* mutant resembled the full *cbf11* deletion in that it showed increased expression of stress-response genes (Fig 4D), and it was resistant to H_2_O_2_ (Fig 3D). Thus, the DNA-binding activity is critical for the Cbf11-mediated repression of stress genes. Overall, these data suggest that Cbf11 transcriptionally regulates some other, stress-unrelated, genes that in turn affect stress-gene expression.

### Multiple lipid-metabolism mutants show derepression of stress genes and resistance to H_2_O_2_

Up to now, we analyzed the *cbf11Δ* response to oxidative stress using several representative genes. To also capture the global picture, we next performed a microarray analysis of gene expression in a timecourse experiment following H_2_O_2_ treatment. This transcriptome analysis confirmed the trends observed so far: many stress-responsive genes were moderately upregulated already in untreated *cbf11Δ* cells and were induced even further upon stress imposition (Fig 4A). Interestingly, the changes in the transcriptome of untreated *cbf11Δ* cells were mostly inductions: 134 genes were ≥2x upregulated, while only 10 genes were ≥2x downregulated compared to untreated WT cells (Fig 4B). This contrasts with the physiological reaction to H_2_O_2_ in WT cells, where stress-gene induction is accompanied by repression of numerous, mostly growth-related genes ((9) and Fig 4B, top panel). Importantly, only 2 out of the 10 genes downregulated in untreated *cbf11Δ* cells are known to be downregulated as part of the core environmental stress response in WT (*car2* and SPBPB7E8.01) (9). These results are in agreement with our earlier notion that the increased stress-gene expression in untreated *cbf11Δ* cells is not merely a consequence of some hypothetical internal oxidative stress activating a genuine stress response.

We previously showed that several genes downregulated in *cbf11Δ* cells are related to lipid metabolism (e.g., FA synthesis) and that Cbf11 directly activates their transcription (11). Indeed, three such direct Cbf11 target genes (*cut6, lcf2, ole1*) were also ≥2x downregulated in untreated *cbf11Δ* cells in the current microarray experiment (Fig 4C). Intriguingly, in *Saccharomyces cerevisiae* decreased FA synthesis caused by inhibition of the acetyl-CoA carboxylase (Cut6 in *S. pombe*) leads to chromatin hyperacetylation and changes in gene expression. Presumably, this is caused by increased availability of acetyl-CoA, which is used as substrate by both ACC and HAT_s_ (17). Lipid-metabolism genes thus represent a potential link between Cbf11 transcription factor activity and changes in stress-gene expression. To examine such potential indirect regulation of stress genes by Cbf11, we assessed stress-gene expression and H_2_O_2_ resistance in other lipid-metabolism mutants. These included the Mga2 transcription factor that also regulates the *ACC/cut6* gene (*mga2Δ;* (12)) and the ACC/Cut6 enzyme itself (*Pcut6MUT* promoter mutant with ~50% reduction in expression (13) and *cut6-621* ts mutant (46)). Strikingly, all mutants showed increased expression of stress genes (Fig 4D), and all but the sick *cut6-621* ts mutant were also resistant to H_2_O_2_ (Fig 4E). Thus our data point to a novel regulatory link between FA synthesis and acetyl-CoA availability on the one hand, and stress-gene expression and cellular resistance to oxidative stress on the other hand.

### Stress genes derepressed in *cbf11Δ* cells show H3K9 hyperacetylation at their promoters

While it was previously shown that decreased FA synthesis leads to chromatin hyperacetylation in *Saccharomyces cerevisiae*, no effect on stress-response genes was reported (17). Intriguingly, we previously showed that the Gcn5/SAGA histone acetyltransferase regulates the expression of stress genes in fission yeast via histone H3 acetylation at lysines 9 and 14 (47, 48). We decided to explore these links further.

First, to determine whether the increased expression of stress genes in the *cbf11Δ* lipid-metabolism mutant was associated with changes in their histone acetylation profiles, we performed ChIP-seq experiments. As previously described (48), the promoters of most stress genes are largely depleted of nucleosomes (compare the H3 levels at promoters and gene bodies in Fig 5A), but we could immunoprecipitate histones both upstream and downstream of the transcription start sites (TSS in Fig 5A). Thus, we determined the occupancy of total histone H3 and H3 acetylated at lysine 9 (H3K9ac), and analyzed their distribution at stress gene bodies and promoter regions. We focused on genes upregulated as part of the core environmental stress response in WT cells (CESR-UP; (9)), and further divided these genes based on their responsiveness to the absence of Cbf11. Strikingly, the CESR-UP genes upregulated in untreated *cbf11Δ* cells (n = 94) showed a marked increase in H3K9 acetylation at their promoters and beginning of gene bodies in *cbf11Δ* cells compared to WT, while no differences in total histone H3 occupancy between the two genotypes were observed (Fig 5A, Fig EV4A). Moreover, the H3K9ac profile of the remaining CESR-UP genes (n = 441) resembled the profile of all fission yeast genes, suggesting specificity of the observed hyperacetylation (Fig 5A, Fig EV4A).

**Figure 5.**
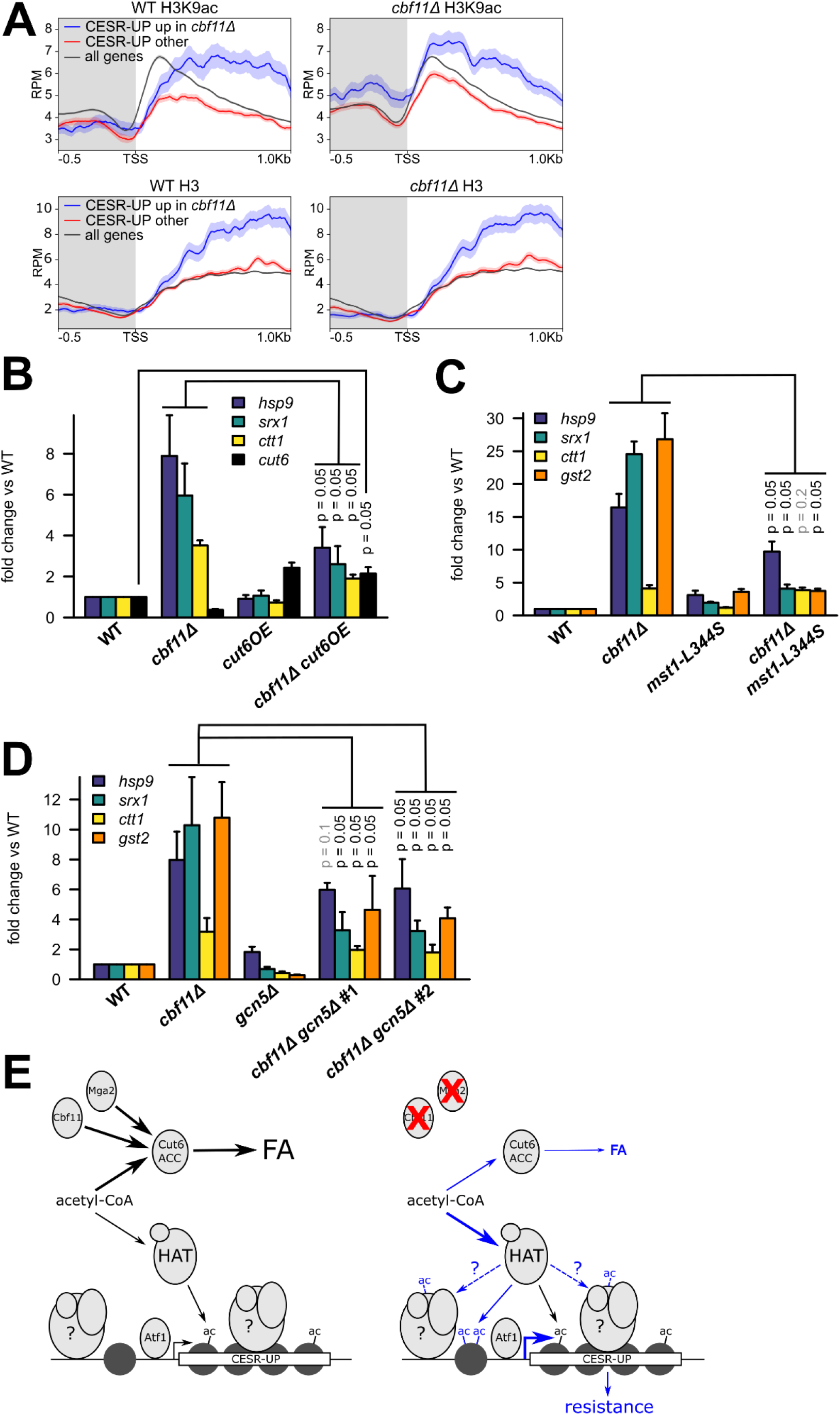
Derepression of stress genes in *cbf11Δ* cells is associated with H3K9 hyperacetylation in their promoters. **(A)** Average gene profiles of total H3 (bottom panels) and acetylated H3K9 (top panels) occupancy at stress gene regions in WT and *cbf11Δ* cells. Genes upregulated as part of the core environmental stress response (CESR-UP, (9)) have been divided into those showing upregulation in untreated *cbf11Δ* cells (blue; n = 94, (11)) and the rest (red; n = 441). Average profile of all fission yeast genes is also shown for comparison (black; n = 6952). The curves represent mean RPM (reads per million mapped reads) values ±SEM; results from one representative biological replicate are shown. The promoter region is shaded. TSS - transcription start site. **(B)** Expression of the indicated stress genes in untreated WT, *cbf11Δ, cut6OE* and *cbf11Δ cut6OE* cells growing in YES medium was analyzed by RT-qPCR. Mean and SD values of three independent replicates are shown. One-sided Mann-Whitney U test was used to determine statistical significance. **(C)** Expression of the indicated stress genes in untreated WT, *cbf11Δ, mst1-L344S* and *cbf11Δ mst1-L344S* cells growing in YES medium was analyzed by RT-qPCR. All cultures were shifted to 32°C (temperature restrictive for the *mst1-L344S* strain) prior to analysis. Mean and SD values of three independent replicates are shown. One-sided Mann-Whitney U test was used to determine statistical significance. **(D)** Expression of the indicated stress genes in untreated WT, *cbf11Δ, gcn5Δ*, and two independent isolates of *cbf11Δ gcn5Δ* cells growing in YES medium was analyzed by RT-qPCR. Mean and SD values of three independent replicates are shown. One-sided Mann-Whitney U test was used to determine statistical significance. **(E)** A model of the crosstalk between lipid metabolism and stress resistance. The left panel shows the WT situation; the right panel shows the situation in mutants with decreased synthesis of FA, with differences highlighted in blue. Hypothetical chromatin remodelers and/or histone modifiers are denoted with a question mark. Dashed lines represent speculative interactions. See Discussion for more details.

Second, we have discovered that even a modest ACC/Cut6 overexpression (~2 fold, Fig 5B) is sufficient to substantially suppress the increased stress-gene expression in the *cbf11Δ* lipid-metabolism mutant (Fig 5B). This observation further confirms the tight relationship between FA metabolism and stress-gene expression. Since ACC activity is hypothesized to affect the general availability of acetyl-CoA (17) we also tested the global levels of acetyl-CoA in lipid metabolism mutants by liquid chromatography-mass spectrometry (LC-MS). While we detected lower acetyl-CoA levels in the *ppc1-537* phosphopantothenate-cysteine ligase mutant strain, which is deficient in CoA synthesis (49), we did not find any significant changes in global acetyl-CoA levels in cell extracts of *cbf11Δ* or *Pcut6MUT* (Fig EV4B). These findings, however, do not rule out the possibility that local, compartmentalized changes in acetyl-CoA availability occur in these two mutants (50), as the nucleocytosolic acetyl-CoA pool, which is directly affected by ACC/Cut6, is not uniform and acetyl-CoA levels at chromatin need not correlate with the global acetyl-CoA levels (51). Moreover, the *S. cerevisiae* AMPK/Snf1 (AMP-activated protein kinase) is a known inhibitor of ACC, and *snf1Δ* budding yeast cells display decreased acetyl-CoA and histone acetylation levels (52). Curiously, the loss of the *S. pombe* AMPK ortholog *ssp2* partially suppressed the stress-gene expression in *cbf11Δ* (Fig EV5A), further stressing the importance for ACC activity and acetyl-CoA availability in regulating the stress-gene expression.

Third, we tested the importance of individual characterized *S. pombe* histone acetyltransferases for the derepression of stress genes in *cbf11Δ* cells. We also included Elp3, the elongator complex acetyltransferase, which was proposed to act only in tRNA modification (53–55), but altered histone acetylation was observed in the *elp3Δ* mutant (56). To this end, we constructed double mutants of *cbf11Δ* and the respective HAT mutations and assayed stress-gene expression in untreated cells using RT-qPCR. We have identified the essential MYST family Mst1 (Fig 5C) and the Gcn5/SAGA acetyltransferases (Fig 5D) as dominant regulators of the stress-gene expression in lipid metabolism mutant cells, while the other tested Gcn5-related N-acetyltransferase family members Hat1 and Elp3 (Fig EV5B), the MYST family protein Mst2 or H3K59-specific HAT Rtt109 (Fig EV5C) were largely dispensable for stress-gene upregulation in untreated *cbf11Δ* cells.

In summary, we have demonstrated a novel regulatory link between FA metabolism and cellular resistance to oxidative stress. When FA synthesis is decreased, a subset of stress-responsive genes becomes derepressed, making cells more resistant to H_2_O_2_. Furthermore, this process is associated with increased histone acetylation at the derepressed stress genes and depends on the activity of specific HATs.

## DISCUSSION

It is well established that metabolism and gene expression are reciprocally regulated to help cells respond efficiently to changes in both intrinsic and extrinsic factors (57). However, the mechanisms underlying this complex regulatory crosstalk, and its diverse implications for cellular physiology are only incompletely understood. In this study, we show that genetic perturbation of lipid metabolism (or more specifically, the biosynthesis of FA) is associated with increased promoter histone H3K9 acetylation and HAT-dependent expression of stress-response genes, which leads to increased resistance of cells to exogenous oxidative stress. Furthermore, both the altered expression of stress genes and increased stress resistance depend on the canonical SAPK (Sty1) pathway.

Both the lipid-metabolism regulators we analyzed (Cbf11, Mga2) and the Sty1 SAPK pathway affect multiple cellular processes, and their mutants show pleiotropic phenotypes. So, how can we distinguish whether our findings represent a genuine functional crosstalk between lipid metabolism and stress resistance or just some indirect effects of cellular stress? First, diverse types of stress activate a common set of stress-response genes and can lead to cross-protection against other, unrelated stresses (9, 58). Conversely, a mild dose of a particular stress can precondition cells to cope with a much higher dose of the same type of stress (59, 60). Several lines of evidence indicate that none of this can explain our observations, and that the increased resistance to oxidative stress of lipid-metabolism mutants is not merely due to the cells being intrinsically stressed. While Sty1 (and partially Atf1) is indeed required for the increased oxidative-stress resistance of lipid-metabolism mutants, we have not detected increased levels of reactive oxygen species in untreated *cbf11Δ* cells (Fig 1D), and neither Sty1 nor Atf1 were hyperactivated in untreated *cbf11Δ* cells (Fig 3A). Moreover, our analysis of the transcriptome of untreated *cbf11Δ* cells identified mainly gene upregulation, which is unlike a typical stress response where a large group of genes is downregulated (Fig 4A, B). Second, the specificity of lipid-metabolism impingement on stress resistance is underlined by the phenotypes of the *cut6* acetyl-CoA carboxylase mutants (Fig 4D, E, Fig 5B). In particular, the well-defined *Pcut6MUT* promoter mutation, which results in ~50% reduction of *cut6* mRNA levels and decreased amount of functional ACC/Cut6 protein without any notable pleiotropic defects (13), does also lead to increased stress-gene expression and resistance to oxidative stress, similar to *cbf11Δ* and *mga2Δ*. Furthermore, the H_2_O_2_-resistant *cbf11Δ* mutant is not resistant to superoxide stress (Fig EV1E), and is actually sensitive to a number of other stresses (12, 39, 40), highlighting the specificity of Cbf11 impact on stress gene expression. Taken together, the crosstalk between lipid metabolism and stress resistance appears to be a genuine phenomenon, and not just a side effect of a pleiotropic mutant phenotype.

How is the crosstalk between lipid metabolism and stress resistance mediated? Our results show that the DNA-binding activity of Cbf11 plays a critical role (Fig 3D), but Cbf11 does not seem to regulate stress-gene expression directly (Fig 3C, Fig EV3). Therefore, some Cbf11 (and/or Mga2) target gene(s) likely provide the connection between lipid metabolism and stress-gene expression. The Cut6 ACC is a strong candidate: 1) the *cut6* gene is regulated both by Cbf11 and Mga2 (12, 13); 2) ACC is a major consumer of acetyl-CoA, capable of affecting histone acetylation levels via limiting acetyl-CoA availability for HAT_s_ (17); 3) hyperactive ACC results in hypoacetylated chromatin and stress-sensitive cells in budding yeast (52); 4) decreased or increased expression of *cut6* alone results in inverse changes in the expression of stress genes (Fig 4D, Fig 5B) and resistance to H_2_O_2_ (Fig 4E); and 5) promoters of stress genes upregulated in the *cbf11Δ* mutant showed increased H3K9 acetylation (Fig 5A, Fig EV4A). Importantly, H3K9 acetylation levels correlate with transcription (61), enhance binding of transcription factors (62) and promote transcriptional elongation by RNA polymerase II (47, 63). Gcn5/SAGA is a major H3K9-targeting HAT in the fission yeast (56), and it is important for proper stress-gene activation during oxidative stress (47). Our data show that Gcn5 is indeed important for the increased stress-gene expression in the *cbf11Δ* lipid mutant (Fig 5D). Strikingly, the evolutionarily conserved MYST family histone acetyltransferase Mst1 (ortholog of human Tip60, *S. cerevisiae* Esa1) with broad enzymatic specificity for both histone and non-histone targets (64, 65) is also strongly required for the increased stress-gene expression in *cbf11Δ* cells (Fig 5C). Since Mst1 is not known to acetylate H3K9, some non-histone proteins, such as chromatin remodelers (66, 67) or other transcription regulators (68), may be regulated by Mst1-dependent acetylation and help project the metabolic state into changes in gene expression (Fig 5E). Interestingly, *S. cerevisiae* Esa1 physically interacts with the stress-responsive transcription factors Msn2, Msn4 and Yap1 (64, 69). Thus, HAT activity (possibly regulated by local acetyl-CoA availability) and stress-gene promoter acetylation represent plausible candidates for the mechanistic link between lipid metabolism and stress-gene expression.

The next question then is how specificity is achieved - why and how only a subset of stress-response genes become specifically upregulated upon perturbation of FA synthesis. Notably, the subset of CESR-UP genes upregulated in untreated *cbf11Δ*, whose promoters show H3K9 hyperacetylation in *cbf11Δ* cells, tend to have above-average nucleosome occupancy in their transcribed regions, unlike the other CESR-UP genes (see the total H3 levels in Fig 5A and Fig EV4A). This suggests that a particular chromatin structure and/or presence of a specific ensemble of chromatin remodelers and histone modifying enzymes make those genes more responsive to changes in the metabolic state. Notably, Gcn5 and Mst1, the HAT_s_ required for increased stress-gene expression in *cbf11Δ* cells, are the catalytic subunits of the histone acetyltransferase modules of transcription co-activator complexes SAGA and NuA4, respectively (70). It is conceivable that other subunits or modules of the SAGA and NuA4 complexes might be responsible for the observed specificity in lipid metabolism-regulated transcription, e.g. by directing HAT complex recruitment to particular genes or by affecting HAT complex interactions with other proteins. Intriguingly, in mammalian cells, lipid-derived acetyl-CoA can provide up to 90% of acetyl-carbon for histone acetylation, and supplementation with the octanoate FA (which is turned into acetyl-CoA by beta-oxidation) results in histone hyperacetylation and induction of specific genes, distinct from those induced by glucose-derived acetyl-CoA (71). Moreover, the ACSS2 acetyl-CoA synthetase was found to be physically associated with chromatin in mouse neurons, where it affects histone acetylation and expression of specific memory-related genes by targeted, on-site production of the acetyl-CoA substrate for HAT_s_ (72). Importantly, in *S. pombe* the Acs1 acetyl-CoA synthetase is the key contributor to the nucleocytosolic acetyl-CoA pool (73), and it localizes predominantly to the nucleus (50). Recruitment of Acs1 to specific genes could thus potentially provide the required specificity in transforming metabolic changes into changes in gene expression.

Finally, what is the purpose of such a crosstalk between lipid metabolism and stress-gene expression? In other words, what is the selective advantage of upregulating specific stress-response genes during decreased FA synthesis? Specific metabolic states are often associated with increased levels of distinct stressors, which can perturb cellular homeostasis. A metabolic control of the stress response would allow cells to better adapt to changes in cellular chemistry and ensure that any potential damage to cellular components is minimized. For example, FA synthesis is downregulated upon carbon-source limitation (74). This is often followed by lipolysis and increased FA oxidation to compensate for the lack of energy resources. Notably, these FA catabolic processes generate increased levels of reactive oxygen species (75). Therefore, a timely mild upregulation of oxidative stress-response genes in response to decreased FA synthesis could represent a useful safety precaution for the cell. Curiously, perturbations of FA metabolism have been linked to altered stress resistance in *Caenorhabditis elegans*, even though the mechanism underlying this connection was not determined (76). Another *C. elegans* study found that the NHR-49 nuclear hormone receptor, a known transcriptional regulator of lipid-metabolism genes, can also (perhaps indirectly) regulate the transcriptional response to fasting and peroxide stress, and is required for resistance to organic peroxides (77). Moreover, an intriguing crosstalk between FA metabolism and stress resistance has been described in mouse. β-hydroxybutyrate, a ketone body produced from oxidized FAs during fasting, prolonged exercise or in patients with diabetes, has been reported to protect against oxidative stress in the mouse kidney. β-hydroxybutyrate inhibits class I histone deacetylases, thereby enhancing H3K9 and H3K14 promoter acetylation and transcription of several stress genes (78). In any case, the proposed mechanism for crosstalk between FA synthesis and resistance to oxidative stress outlined in Fig 5E needs to be tested experimentally in future studies.

## DATA AVAILABILITY

The datasets produced in this study are available from the ArrayExpress database (https://www.ebi.ac.uk/arrayexpress/) under accession numbers E-MTAB-6761 (expression microarrays) and E-MTAB-11081 (ChIP-seq).

The scripts used for ChIP-seq data processing and analysis are available from https://github.com/mprevorovsky/ox-stress_histones.

## ACKNOWLEDGEMENTS

The authors would like to thank Eishi and Chiaki Noguchi for providing the *mst1ts* strain, Anna Janovská and the Laboratory of Mass Spectrometry at Biocev Research Center, Faculty of Science, Charles University for performing the LC-MS analyses, NBRP Japan for providing the *ppc1-537* strain, Simona Veselá, Patrik Hohoš and Viacheslav Zemlianski for help with the *ssp2Δ* and *cbf11Δ ssp2Δ* strain construction, all members of the GenoMik and ReGenEx groups for their support and insightful discussions, and Eva Krellerová, Adéla Kmochová and Kateřina Jelínková for their technical assistance.

## AUTHOR CONTRIBUTIONS

Conceptualized the study and designed the experiments: MP, EH, JA, JP, JB

Performed the experiments: JP, CS-C, PD, AM, LdC, MP Analyzed the data: JP, CS-C, PD, MP, AM, LdC, EH, JA

Wrote the manuscript: MP, JP, CS-C, EH, JA. All the other authors contributed to the final revised version.

