## Supplemental Data for "Perturbed fatty-acid metabolism is linked to localized chromatin hyperacetylation, increased stress-response gene expression and resistance to oxidative stress"

### EXPANDED VIEW

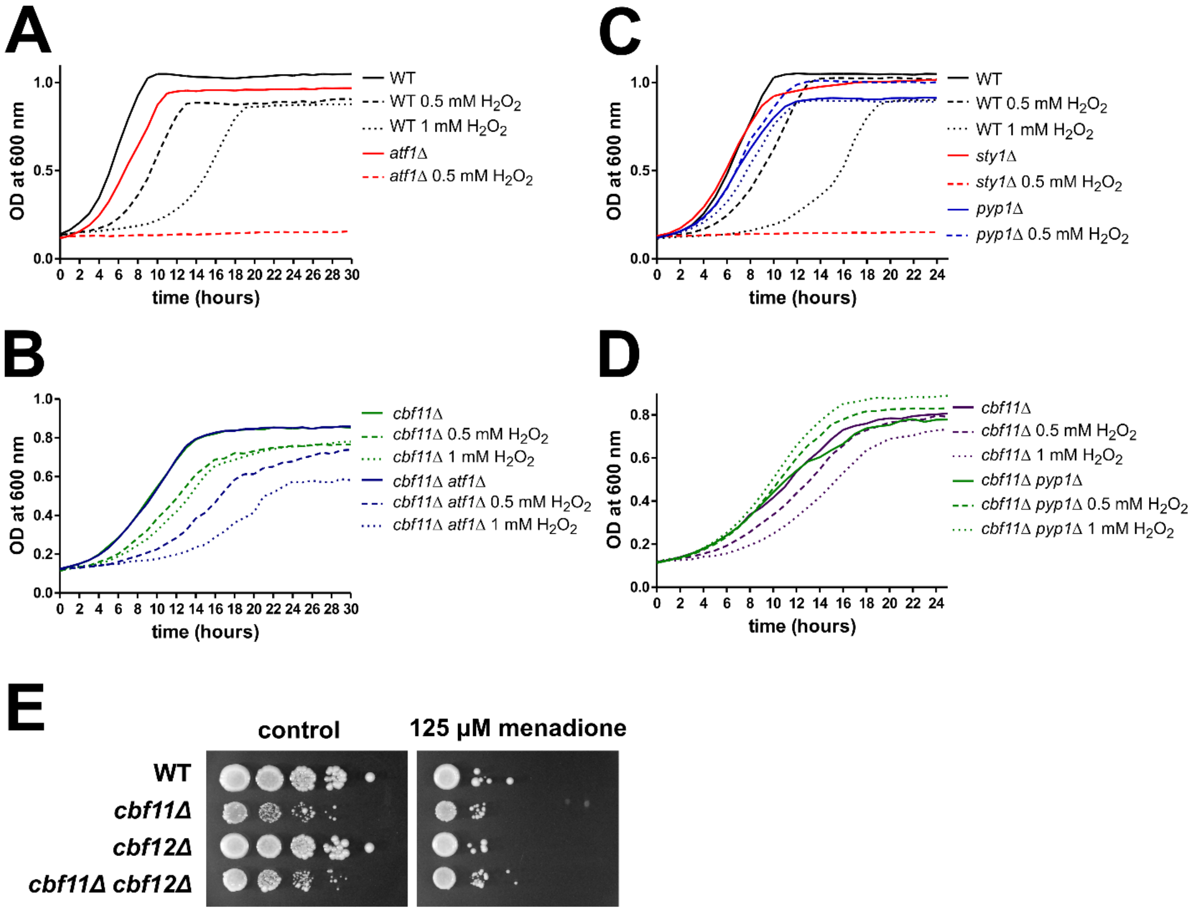


**Figure EV1 - Resistance of *cbf11Δ* cells to oxidative stress partially depends on Atf1 and is specific for hydrogen peroxide.**

**(A-D)** Growth curves of stress-related mutants in the presence or absence of the indicated concentrations of $H_2O_2$ in YES medium. The *pyp1Δ* and *sty1Δ* strains represent strongly resistant and strongly sensitive controls, respectively. **(E)** Survival and growth under superoxide stress of WT, *cbf11Δ*, *cbf12Δ* and *cbf11Δ cbf12Δ* cultures spotted on YES plates containing 125 μM menadione.

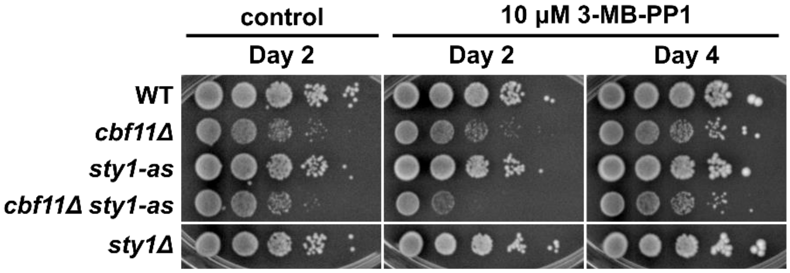


**Figure EV2 - Lack of Sty1 activity further impairs growth of *cbf11Δ* cells.**
Exponentially growing WT, *cbf11Δ*, *sty1-as*, *cbf11Δ sty1-as*, and *sty1Δ* cultures were spotted on YES plates containing 10 µM Sty1-as inhibitor 3-MB-PP1 and incubated for the indicated number of days.

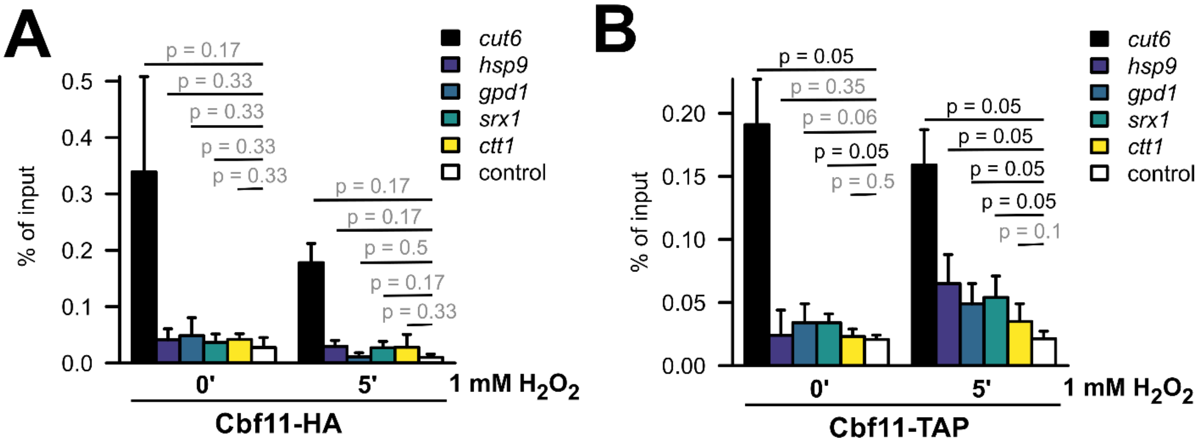


**Figure EV3 - Cbf11 does not bind to Atf1 binding sites at stress-gene promoters.**
**(A, B)** Recruitment of Cbf11-HA and Cbf11-TAP, respectively, to known Atf1 binding sites in the indicated stress gene promoters was analyzed by ChIP-qPCR in cells treated or not with 1 mM $H_2O_2$ for 5 min in EMM medium. The *cut6* promoter is a positive control for Cbf11 binding (13); ”control” is a locus with no expected Cbf11 binding. Mean and SD values of two (A) and three (B) independent replicates are shown. One-sided Mann-Whitney U test was used to determine statistical significance.

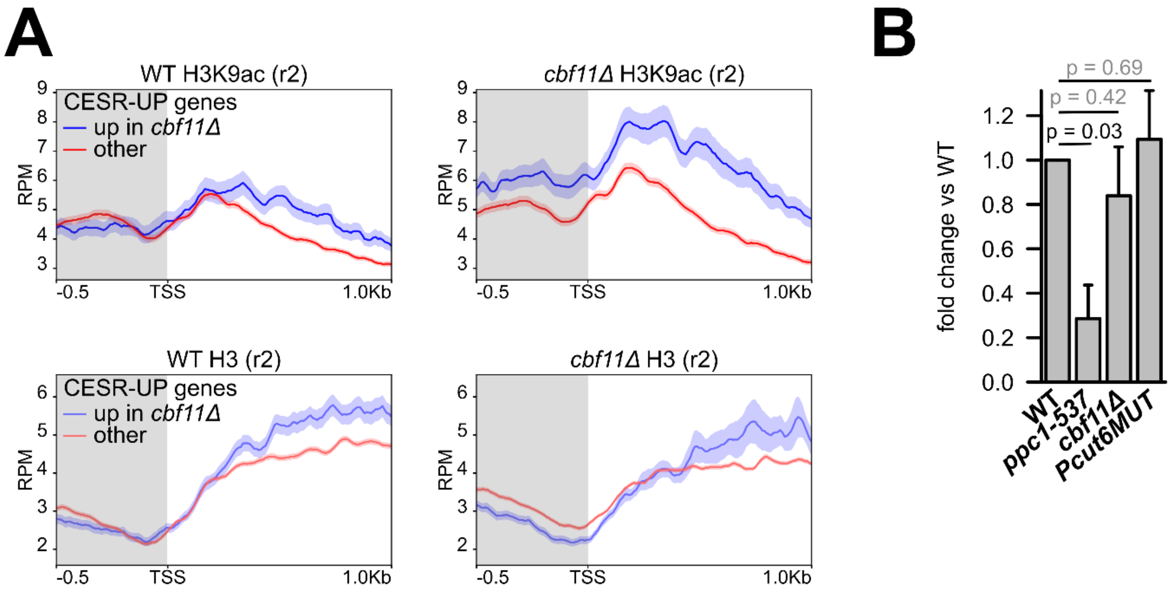


**Figure EV4 - Derepression of stress genes in *cbf11Δ* cells is associated with H3K9 hyperacetylation in their promoters (the second replicate) but not with altered total cellular acetyl-CoA levels.**

**(A)** Average gene profiles of total H3 (bottom panels) and acetylated H3K9 (top panels) occupancy at stress-gene regions in WT and *cbf11Δ* cells, respectively. Genes upregulated as part of the core environmental stress response (CESR-UP, (9)) have been divided into those showing upregulation in untreated *cbf11Δ* cells (blue; n = 94, (11)) and the rest (red; n = 441). Average profile of all fission yeast genes is also shown for comparison (black; n = 6952). The curves represent mean RPM (reads per million mapped reads) values ± SEM. The promoter region is shaded. TSS - transcription start site.

**(B)** Total cellular acetyl-CoA levels in WT, *ppc1-537*, *cbf11Δ* and *Pcut6MUT* cell extracts were determined by LC-MS. Mean and SD values of four independent replicates for *ppc1-537* and five independent replicates for all other strains tested are shown. Two-sided Mann-Whitney U test was used to determine statistical significance.

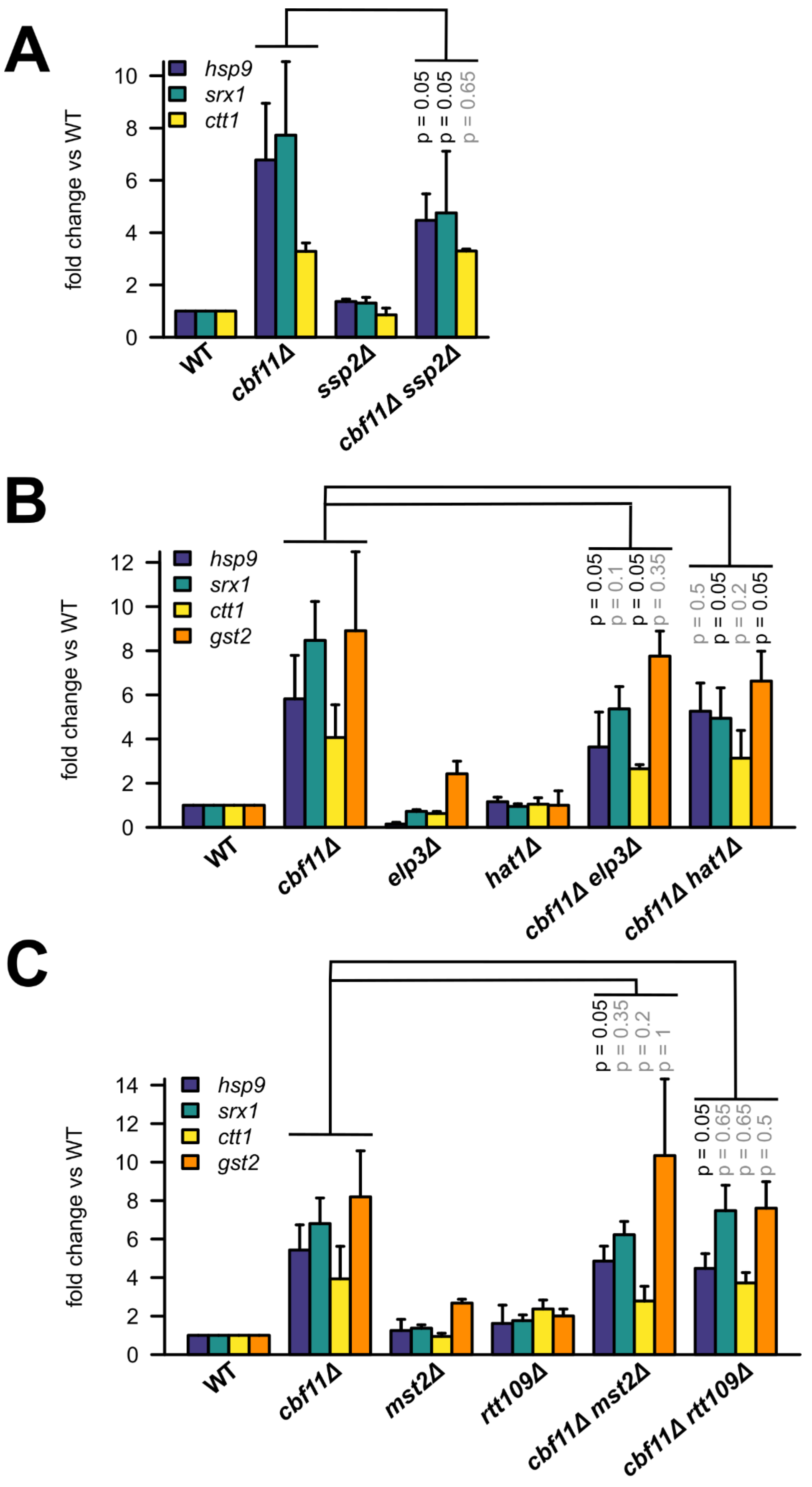
A
fold change vs WT
hsp9
srx1
ctt1
p = 0.05
p = 0.05
p = 0.65
WT
cbf11Δ
ssp2Δ
cbf11Δ ssp2Δ
B
fold change vs WT
hsp9
srx1
ctt1
gst2
p = 0.05
p = 0.1
p = 0.05
p = 0.35
p = 0.5
p = 0.05
p = 0.2
p = 0.05
WT
cbf11Δ
elp3Δ
hat1Δ
cbf11Δ elp3Δ
cbf11Δ hat1Δ
C
fold change vs WT
hsp9
srx1
ctt1
gst2
p = 0.05
p = 0.35
p = 0.2
p = 1
p = 0.05
p = 0.65
p = 0.65
p = 0.5
WT
cbf11Δ
mst2Δ
rtt109Δ
cbf11Δ mst2Δ
cbf11Δ rtt109Δ

**Figure EV5 - AMPK and histone acetyltransferases Elp3, Hat1, Rtt109 and Mst2 have limited effect on stress gene derepression in *cbf11Δ* cells.**

**(A, B, C)** Expression of the indicated stress genes in cells growing in YES medium was analyzed by RT-qPCR. Mean and SD values of three independent replicates are shown. One-sided Mann-Whitney U test was used to determine statistical significance.

### APPENDIX

### Supplementary Table 1 – List of oligonucleotides

| ID | Sequence | Experiment |
|---|---|---|
| MP137 | TCCTCATGCTATCATGCGTCTT | RT-qPCR *act1* (fwd) |
| MP138 | CCACGCTCCATGAGAATCTTC | RT-qPCR *act1* (rev) |
| MP169 | GGTCTATGTTCCCACTGTTT | RT-qPCR *rho1* (fwd) |
| MP170 | CTTCTTGTCCAGCCGTA | RT-qPCR *rho1* (rev) |
| MaP173 | CCGACCCTTCATTAACATTAC | RT-qPCR *atf1* (fwd) |
| MaP174 | CCGTTGGCATATCAGAATTAG | RT-qPCR *atf1* (rev) |
| PD01 | CCGACGACATTACCAAGTA | RT-qPCR *ctt1* (fwd) |
| PD02 | ACGAGCAGTATCAGGAGTA | RT-qPCR *ctt1* (rev) |
| PD05 | TGCCTTCTTATCCCTCCT | RT-qPCR *pyp2* (fwd) |
| PD06 | ACGACGTTGCTGGATTTA | RT-qPCR *pyp2* (rev) |
| AJ33 | aaaatctagaGGGAACGTGTCAAAGATGCC | *mga2* KO cloning primer, upstream/outside, fwd (XbaI) |
| AJ34 | aaaactcgagTTGAGAGTACTTTGAAATGCAATAGA | *mga2* KO cloning primer, upstream/inside, rev (XhoI), product size 738 bp (with AJ33) |
| AJ35 | aaaatctagaGCAGATCGCATACCGGTTCA | *mga2* KO cloning primer, downstream/outside, rev (XbaI) |
| AJ36 | aaaaagatctTTTGACGAACAACCATGC | *mga2* KO cloning primer, downstream/inside, fwd (BglII), product size 424 bp (with AJ35) |
| AJ37 | TCTGTTGGCCGGAATTTGG | *mga2* KO checking primer, upstream, fwd, product size 983 bp (with MP144) |
| AJ38 | ACGTCCTCTCCCATGCTTGA | *mga2* KO checking primer, downstream, rev, product size 967 (with MP33) |
| MaP90 | GTATCGTCTTGCTCGGTT | ChIP-qPCR *atf1* (fwd) |
| MaP91 | CCACACTTCCACCTGTTT | ChIP-qPCR *atf1* (rev) |
| PD03 | GAATTACCAACGTCATATTTGC | ChIP-qPCR *ctt1* (fwd) |
| PD04 | ACTACGATAGGCTGTAGAAGA | ChIP-qPCR *ctt1* (rev) |
| MaP96 | TTGCTACAGGAAGAGGAAG | ChIP-qPCR *cut6* (fwd) |
| MaP97 | TAGAAAAGTTGGATGCGTG | ChIP-qPCR *cut6* (rev) |
| PD07 | GCGTCACTCGTCACATTA | ChIP-qPCR *pyp2* (fwd) |
| PD08 | TGCTAAGCGACCGTTTATT | ChIP-qPCR *pyp2* (rev) |
| MP88 | AGCTGCTAGACACCTTCAAA | ChIP-qPCR intergenic locus P4 (fwd) |
| MP89 | CCTACGGTCAAGAGAAAACT | ChIP-qPCR intergenic locus P4 (rev) |
| MP33 | GCGCACGTCAAGACTGTC | universal KO checking primer (*kanMX6/natMX6* cassette), downstream, fwd |

| | | |
|---|---|---|
| MP144 | GTCGTTAGAACGCGGCTACA | universal KO checking primer, upstream, rev |
| MP150 | AGGGATCGAAAGACATCCGC | *cbf11* KO checking primer, upstream, fwd, product size 953 bp (with MP144) |
| MP151 | GCTTGTACACACGGCCTTCAA | *cbf11* KO checking primer, downstream, rev, product size 823 (with MP33) |
| MaP169 | AAAAAGCTAAATGATGCT | TAP tag |
| MP28 | GATACAGCAACTCCTCCCG | *cbf11* KO checking primer, outer genomic sequence, rev |
| MP53 | AGGTTAATACGCAATGG | *cbf11* ORF, to amplify region around DBM (with MP54), rev |
| MP54 | TTAGTATTGTCTCCAAACC | *cbf11* ORF, to amplify region around DBM (with MP53), fwd |
| AJ11 | GCGCAGCTCAGTTTTAGAGCTAGAAATAGCAAGTTAAAATAA | CRISPR/Cas9, first half of sgRNA against *natMX6* (TEF promoter), overlap into pMZ374, fwd |
| AJ12 | ACGTCAAGACTTCTTCGGTACAGGTTATGTTTTTTGGCAACA | CRISPR/Cas9, second half of sgRNA against *natMX6* (TEF promoter), overlap into pMZ374, rev |
| AJ29 | cctttataacGTTTTAGAGCTAGAAATAGCAAGTTAAAATAA | CRISPR/Cas9, first half of sgRNA against *cbf11* ORF (next to desired DBM), overlap into pMZ374, fwd |
| AJ30 | ctgacgactgTTCTTCGGTACAGGTTATGTTTTTTGGCAACA | CRISPR/Cas9, second half of sgRNA against *cbf11* ORF (next to desired DBM), overlap into pMZ374, rev |
| JT68 | CAAGGCCAAGGAATCCATTA | RT-qPCR *hsp9* ORF (fwd) |
| JT69 | GAGCCTTGTCATGAGCCTCT | RT-qPCR *hsp9* ORF (rev) |
| JT70 | TTCATGCGGTTTGACTTCAG | RT-qPCR *srx1* ORF (fwd) |
| JT71 | CCCCCAAAGGCAAAATAATA | RT-qPCR *srx1* ORF (rev) |

**Supplementary Table 2 – List of plasmids**

| ID | Vector | Experiment/use | Source |
|---|---|---|---|
| pMP90 | pCloneNAT1 | KO vector for *cbf11* (*kanR* cassette) | This study |
| pMP91 | pCloneNAT1 | KO vector for *cbf11* (*natR* cassette) | Ref. 1 |
| pMP148 | pCloneNAT1 | KO vector for *ssp2* (*natR* cassette) | This study |
| pMP161 | pCloneNAT1 | KO vector for *mga2* (*natR* cassette) | This study |
| pMP134 | pMZ374 | Cas9/sgRNA_TEFp; Cas9, sgRNA against *natMX6* (TEF promoter) | This study |
| pMP153 | pMZ374 | Cas9/sgRNA_cbf11; Cas9, sgRNA against *cbf11* ORF (next to desired DBM) | This study |
| pMZ374 | | Cas9/sgRNA_empty; Cas9, no sgRNA | Addgene plasmid # 59896 / Mikel Zaratiegui |
| pMaP11 | pUC19 | *cbf11DBM-TAP*, template for homologous recombination | This study |
| pMaP27 | pUC19 | *cbf11-TAP*, template for homologous recombination | This study |
| p407.C169S | pREP | fluorescence measurement of intracellular $H_2O_2$ | Ref. 2 |

### Supplementary Table 3 – List of strains

| ID | Genotype | Source | Figure # |
|---|---|---|---|
| JB32 | *h+* | Lab stock | Fig 1B bottom panel, Fig 1C, Fig 2B, Fig 2C, Fig 3D, Fig 4A, Fig 4B, Fig 4D, Fig 4E, Fig 5A-D, Fig EV2, Fig EV4 |
| MP114 | *h+ cbf11Δ::natR* | Ref. 1 | Fig 1C, Fig 2C |
| MP21 | *h+ cbf12Δ::natR* | Ref. 1 | Fig 1C, Fig 4A |
| MP22 | *h+ cbf11Δ::kanR* | Ref. 1 | Fig 4A, Fig4B |
| MP25 | *h+ cbf11Δ::kanR cbf12Δ::natR* | Ref. 1 | Fig 1C |
| JB146 | *h- atf1Δ::ura4 ura4-D18* | Lab stock | Fig 2B, Fig 2C |
| MP367 | *h- ura4-D18 atf1Δ::ura4 cbf11Δ::natR* | This study | Fig 2B, Fig 2C |
| MP44 | *h+ cbf11Δ::kanR* | Ref. 1 | Fig 1B bottom panel, Fig 2B, Fig 3D, Fig 4E, Fig 5A, Fig 5B, Fig EV2, Fig EV4 |
| JB149 | *h- sty1Δ::ura4 ura4-D18* | Lab stock | Fig 2B, Fig EV2 |
| MP809 | *h+ sty1.T97A ura4-D18* | Ref. 3 | Fig 2A, Fig EV2 |
| MP810 | *h+ sty1.T97A ura4-D18 Δcbf11::natR* | This study | Fig 2A, Fig EV2 |
| MP705 | *h+ cbf11-ctap4* | This study | Fig 3D |
| MP712 | *h+ cbf11DBM-ctap4* | This study | Fig 3D, Fig 4D, Fig 4E |
| MP815 | *h+ mga2Δ::natR* | This study | Fig 4D, Fig 4E |
| MP636 | *h- Pcut6MUT* | This study | Fig 4D, Fig 4E |
| MP218 | *h+ cut6-621* | Ref. 4 | Fig 4D, Fig 4E |
| MP19 | *h+ cbf11-ctap4::natR* | Ref. 1 | Fig 3C |
| MP15 | *h- cbf11-ctap4::natR ura4-D18 leu1-32 ade6-M216* | Ref. 1 | |
| MaP70 | *h- cbf11-3HA::natMX6 ura4-D18 leu1-32 ade6-M216* | This study | |
| MP670 | *h- cbf11DBM-3HA::nonfunctional_natMX6 ura4-D18 leu1? ade6?* | This study | |
| MP865 (MS112) | *h+ gcn5::kanMX6* | Ref. 5 | Fig 5D |
| MP879 | *h+ gcn5::kanMX6 Δcbf11::natR* | This study | Fig 5D |
| MP880 | *h+ gcn5::kanMX6 Δcbf11::natR* | This study | Fig 5D |
| 972 | *h-* | Lab stock | Fig 1B top panel, Fig 1F, Fig 3A |
| AV18 | *h- sty1Δ::kanMX6* | Ref. 6 | Fig 3A |
| CS38 | *h- atf1-HA:natMX6* | Ref. 7 | Fig 3B |
| CS89 | *h- cbf11Δ::kanMX6* | This study | Fig 1B top panel, Fig 3A |

| | | | |
|---|---|---|---|
| CS95 | *h? cbf11Δ::kanMX6 atf1Δ::natMX6* | This study | Fig EV1B |
| CS99 | *h+ cbf11Δ::kanMX6* | This study | Fig EV1B |
| CS102 | *h- cbf11-HA:kanMX6* | This study | Fig EV3A |
| CS110 | *h- cbf12Δ::kanMX6* | This study | Fig 1F |
| CS129 | *h+ cbf11-TAP:kanMX6* | This study | Fig EV3B |
| CS142 | *h? atf1-HA:natMX6 cbf11::kanMX6* | This study | Fig 3B |
| MS98 | *h- atf1Δ::natMX6* | Ref. 8 | Fig EV1A |
| HM123 | *h- leu1-32* | Lab stock | |
| MP895 (IV16) | *h+ elp3::natMX6* | Ref. 8 | Fig EV5B |
| MP904 (ENY2778) | *h- mst1-L344S-5FLAG:kanMX6 leu1-32 ura4-D18* | Ref. 9 | Fig 5C |
| MP905 (IV65) | *h- hat1::kanMX6* | Lab stock | Fig EV5B |
| MP907 (JA2344) | *h+ mst2::kanR* | Lab stock | Fig EV5C |
| MP908 (JA2242) | *h- rtt109:kanR* | Lab stock | Fig EV5C |
| MP925 | *h+ elp3::natMX6 cbf11::kanR* | This study | Fig EV5B |
| MP927 | *h- hat1::kanMX6 cbf11::natR* | This study | Fig EV5B |
| MP948 | *h- mst1-L3444S-5FLAG:kanMX6 leu1-32 ura4-D18 cbf11::natR* | This study | Fig 5C |
| MP950 | *h+ mst2::kanR cbf11::natR* | This study | Fig EV5C |
| MP951 | *h- rtt109:kanR cbf11::natR* | This study | Fig EV5C |
| MP550 | *h+s natMX6-Padh1-3HA:cut6+* | Ref. 10 | Fig 5B |
| MP555 | *h+ Δcbf11::kanR natMX6-Padh1-3HA:cut6+* | Ref. 10 | Fig 5B |
| MP635 | *h+ Δssp2::natR* | This study | Fig EV5A |
| MP855 | *h+ Δssp2::natR Δcbf11::hygR* | This study | Fig EV5A |
| MP606 | *h- ppc1-537* | Ref. 11 | Fig EV4B |
